# An ABM of biting midge dynamics to understand Bluetongue outbreaks

**DOI:** 10.1101/2022.09.26.509502

**Authors:** Shane L. Gladson, Tracy L. Stepien

## Abstract

Bluetongue (BT) is a well-known vector-borne disease that infects ruminants such as sheep, cattle, and deer with high mortality rates. Recent outbreaks in Europe highlight the importance of understanding vector-host dynamics and potential courses of action to mitigate the damage that can be done by BT. We present an agent-based model (ABM), entitled MidgePy, that focuses on the movement of individual *Culicoides* spp. biting midges and their interactions with ruminants to understand their role as vectors in BT outbreaks, especially in regions that do not regularly experience outbreaks. Sensitivity analysis is performed and results indicate that midge survival rate has a significant impact on the probability of a BTV outbreak as well as its severity. Parameter regions where outbreaks are more likely to occur are determined, with an increase in environmental temperature corresponding with an increased probability of outbreak, where midge flight activity is used as a proxy for temperature. This suggests that future methods to control BT spread could combine large-scale vaccination programs with biting midge population control measures such as the use of pesticides. Spatial heterogeneity in the environment is also explored to give insight on optimal farm layouts to reduce the potential for BT outbreaks.

## 1 Introduction

In the late 2000s and early 2010s, multiple Bluetongue (BT) disease outbreaks occurred across Northern Europe, a region which did not previously witness BT spread. The resulting epizootic, fueled by an abundance of immunologically naive livestock, lasted multiple years until it was finally brought under control through mass vaccination programs and the restriction of livestock movement across country borders (Gethmann et al. [2020]). The epizootic had a significant economic impact on the livestock industry, and its persistence represents a continued threat to the European agricultural economy (Conraths et al. [2009]).

Occasional epizootics occur in any given year now and result in losses amounting to hundreds of millions of dollars due to necessary vaccination programs or lost revenue (Mayo et al. [2020]). Furthermore, livestock infected with BT experience high morbidity and mortality rates (Saegerman et al. [2008]). Understanding how BT arrived in Northern Europe is of interest to help prevent or mitigate potential outbreaks in new regions. In fact, BT is now considered a prime example of a disease whose range is changing due to climate change bringing warmer temperatures farther north (Brand and Keeling [2017], Samy and Peterson [2016]).

BT is a vector-borne disease caused by Bluetongue Virus (BTV) that is transmitted by *Culicoides* spp. biting midges (Diptera: Ceratopogonidae) and affects a wide range of ruminants such as sheep, cattle, and deer. Thus, the dispersal of *Culicoides* spp. biting midges plays an integral role in the spreading of BT between farms which are geographically distant (Pedgley and Brooksby [1983], Alba et al. [2004], Sellers and Maarouf 1990, 1989], Chapman et al. [2010]).

Previous mathematical models have focused on various aspects of the spread of BT among animal populations with a considerable focus on the Northern Europe BTV outbreaks of the 2000–2010s (see the review paper by Courtejoie et al. [2018] and the references therein). For example, Gubbins et al. [2008] performed sensitivity analysis on a temperature-dependent model considering a population with two host species to predict the risk of BT in Great Britain. Szmaragd et al. [2009, 2010] designed a model that examined the effects of vaccine uptake among farms in Great Britain following the 2006 BTV-8 outbreak. Gourley et al. [2011] used a delay differential equation model to investigate reproduction numbers for BT. Using historical wind data, Sedda et al. [2012] retroactively simulated the spread of the 2006 BTV outbreak in Northern Europe by considering a spatio-temporal environment. Guis et al. [2012] developed a climate driven model, which they used to predict temporal changes in *R*_0_, and subsequently predict an increase in the future risk of BT across Northern Europe. Additionally, Turner et al. [2012] modeled farm–to–farm BT spread across Eastern England, which also considered the seasonal effect on transmission rate. A model that incorporated a temperature-dependent incubation period for BT was designed by Li and Zhao [2019] to predict the reproduction ratio and perform sensitivity analysis.

However, there is a need to examine the potential of BT spread and its impact in other locations that have not been historically affected as substantially as Northern Europe. In particular, while BTV is transmitted in many parts of North America, there exists only one confirmed vector: *C. sonorensis* (Gerry et al. [2001]). This poses a conundrum since the primary range for *C. sonorensis* does not extend to the Southeastern United States, yet there have been confirmed cases of BT in that region (Smith and Stallknecht [1996]). Entomologists have identified potential BTV vector species in the Southeastern United States including *C. stellifer, C. debilipalpis*, and *C. pallidicornis*, however transmission of BT by these species has yet to be directly observed (McGregor et al. [2019]).

Here, we develop a mathematical model to study the potential of BT outbreaks on a single farm in regions not regularly experiencing outbreaks that incorporates the movement of midges and their interactions with ruminants. To include a spatial domain, we develop an agent-based model (ABM) that allows for much finer resolution in the domain than a multi-patch ordinary differential equations model without the necessity of developing a partial differential equations model. The use of agent-based models (ABMs) has been well-established for understanding dynamics of disease spread and the navigation of insects. For example, Smith et al. [2018] published a comprehensive review on the use of ABMs to model malaria transmission. These models were each designed specifically to either understand the role of environment heterogeneity in malaria spread, analyze the effectiveness of intervention strategies, or for parameter estimation. The flight patterns of insects such as moths (Bau and Cardé [2015], Liberzon et al. [2018], Stepien et al. [2020], Golov et al. [2021]), locusts (Topaz et al. [2008], Bernoff et al. [2020]), flies (Lin et al. [2015], Alderton et al. [2018], Leitch et al. [2021], Diouf et al. [2022]), honeybees (Dorin et al. [2022]), and butterflies (Grant et al. [2018]) have all been studied with ABMs using a variety of modes including simple random walks and directed movement toward a target.

This paper aims to understand the impacts of midge movement on the transmission of BT using an ABM, and how aspects such as the number of initial infected midges, the daily survival rate of the midges, the extrinsic incubation period of midges inoculated with BTV, and the probability of BTV transmission between vector and host can affect the probability of an outbreak on a single farm. We also study the effects of temperature using the number of actively flying midges in a simulation as a proxy to determine parameter regions where outbreaks are more likely. Spatial heterogeneity in the environment is explored to give insight on optimal farm layouts to reduce the potential for BT outbreaks. While there exist many parameters related to both vectors and hosts which are known to affect the spread of BT, this paper focuses on those associated with midges.

The outline of this paper is as follows: in Section 2, we provide biological information on BT and its spread. In Section 3, we describe an agent-based model, entitled MidgePy, that examines the spread of BT on a small-scale large mammal farm of approximately one square kilometer. In Section 4, we perform sensitivity analysis of *Culicoides* spp. survival rates, the extrinsic incubation period of BTV, and BT transmission probability, as well as an analysis on outbreak probability and the effects of a heterogeneous spatial domain. Finally, in Section 5, we summarize our results including their application to real-world efforts to combat BT outbreaks.

## 2 Bluetongue (BT)

Bluetongue virus (BTV) is a virus from the family *Reoviridae*, genus *Orbivirus* (Rivera et al. [2021]). *BTV causes Bluetongue (BT), a hemorrhagic disease in ruminants that is transmitted by many species of Culicoides* spp. biting midges (Mellor [1990], Mellor et al. [2000], Mellor [2000]). There are more than 20 known serotypes of BTV that circulate between different regions including the Middle East, Europe, and North America (Gerbier et al. [2008], Maclachlan et al. [2015], Saegerman et al. [2008]).

While vaccines exist that prevent BT, the virus is not immunologically simple and thus the vaccines must be serotype specific. With the large number of serotypes, each with varying virulence, it is difficult or impossible to vaccinate livestock against all strains (Noad and Roy [2009]). In addition, recent studies have shown that, due to the segmented genome of BTV, the use of live attenuated viruses to vaccinate ruminants has the possibility of introducing new genetic material to environments, increasing the risk of creating new BTV serotypes (Rojas et al. [2021]).

Transmission begins when a competent *Culicoides* spp. vector bites a viremic ruminant, ingesting blood that contains BTV. The virus then undergoes an incubation period inside the midge, during which it goes through multiple stages in the incubation cycle before eventually replicating within the midge and reaching its salivary glands (Mellor [2000], Mellor et al. [2000]). The length of this process is known as the Extrinsic Incubation Period (EIP) (Carpenter et al. [2011]). In *Culicoides* spp. vectors, this process typically lasts 14 days, however, it is significantly affected by the ambient temperature, which is accompanied by increasing *Culicoides* spp. activity (Tsutsui et al. [2011], Mayo et al. [2020], Tugwell et al. [2021]). Additionally, the biting rate in *Culicoides* spp. has been shown to be positively correlated with disease transmission rates (Elbers and Meiswinkel [2014]). Though it has recently been shown that horizontal transmission of BT between ruminants is possible (Maclachlan et al. [2019]), the goal of this paper is to focus solely on the vector-host transmission cycle, therefore, horizontal transmission of BT will not be considered here.

## 3 Methods

We present a spatially and temporally explicit agent-based model (ABM), entitled MidgePy, that characterizes the actions of ruminants and *Culicoides* spp. biting midges and the spread of Bluetongue (BT) among these populations. The spatial landscape of our study is based on a cervid farm in North Florida that has been the focus of previous BT dynamics studies (McGregor et al. [2019], Erram et al. [2019], Erram and Burkett-Cadena [2018]). However, the spatial landscape in MidgePy is customizable so that the outbreak likelihood given the introduction of BTV-infected midges to any region may be examined.

The MidgePy algorithm is depicted as a flow chart in Fig. 1, which we subsequently describe.

**Figure 1.**
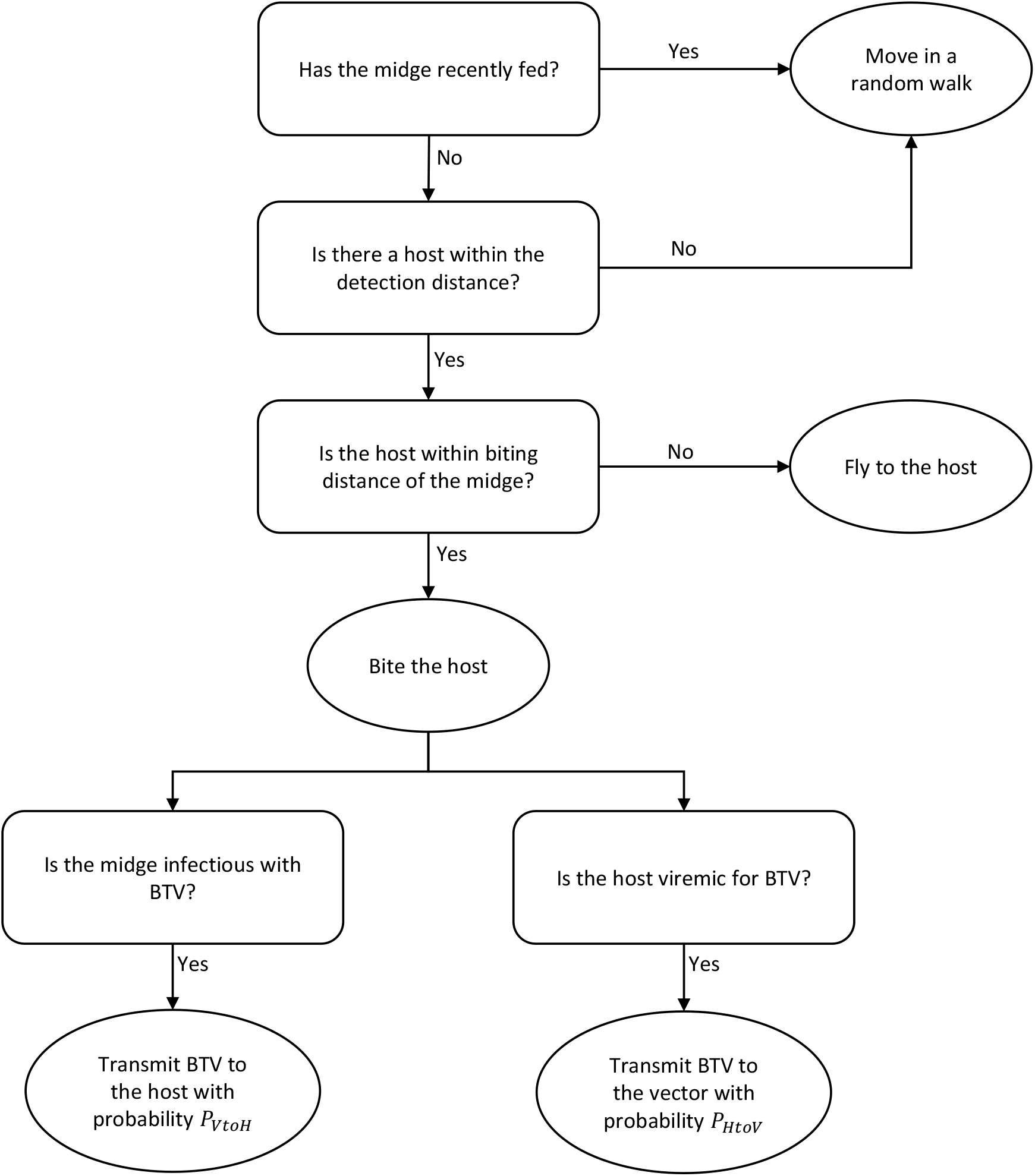
Flowchart of the MidgePy algorithm that determines *Culicoides* spp. behavior at each step.

### 3.1 Model State Variables

MidgePy is comprised of two main classes of agents: a midge swarm class, named MidgeSwarm, and a ruminant swarm class, named HostSwarm. Each class holds arrays containing information on each agent, such as their location and whether or not they are infected with BT. The length of the array indicates the population size of the respective agent type. In particular, the MidgeSwarm and HostSwarm classes contain arrays of lengths *γP*_*d*_ and *P*_*d*_, respectively, where *γ* : 1 is the ratio of midges to ruminants in a simulation. See Table 1 for a list of model parameters and their values.

**Table 1:** Model parameters of the agent-based model MidgePy. For all parameter values that do not have a reference, see Section 3.

| Parameter | Description | Value | Units | Reference |
| --- | --- | --- | --- | --- |
| $\Delta t$ | Time step | 60 | s | |
| $T_{\text{full}}$ | Length of time after a blood meal that a midge remains not hungry | 2 | days | |
| $T_d$ | Length of each day simulated | 5 | hours | Fall et al. [2015], González et al. [2017], Blackwell [1997] |
| $P_d$ | Ruminant population size | 100 | | |
| $\gamma$ | Number of midges per ruminant | 100 : 1 | | |
| $v_r$ | Flight speed of a roaming midge | 0.13 | m/s | Sedda et al. [2012] |
| $v_f$ | Flight speed of an active midge | 0.5 | m/s | Sedda et al. [2012], Gethmann et al. [2020] |
| $d_{\text{det}}$ | Maximum distance a midge can detect a ruminant | 300 | m | |
| $d_{\text{bite}}$ | Minimum distance required for a midge to bite a ruminant | $v_f \Delta t$ | m | |
| $P_{\text{VtoH}}$ | Probability of BTV transmission from vector to host | 0.9 | | Bessell et al. [2016] |
| $P_{\text{HtoV}}$ | Probability of BTV transmission from host to vector | 0.14 | | Carpenter et al. [2013] |
| $\rho_H$ | Extrinsic incubation period of BTV in ruminants | 2 | days | |
| $\rho$ | Extrinsic incubation period of BTV in *Culicoides* spp. | 10–20 | days | Tsutsui et al. [2011], Mayo et al. [2020], Tugwell et al. [2021], Wittmann et al. [2002] |
| $\alpha$ | *Culicoides* spp. daily survival probability | 0.5–0.9 | | Wittmann et al. [2002], Lysyk and Danyk [2007], Gerry and Mullen [2000] |
| $I_0$ | Number of midges infected with BTV at initialization | 1–5 | | |

An increase in the midges-per-ruminant ratio *γ* significantly increased the computational cost when running the model, and so we ran simulations to determine a value of *γ* that would allow for a reasonable computational time yet be representative of the full population dynamics. Initial simulations were run setting *γ* = 30, and then increased to *γ* = 100. Simulations run with *γ >* 100 showed no significant differences in model output. Therefore it was decided that *γ* = 100 was a suitable parameter value for this analysis.

### 3.2 Initialization of Simulations

We considered both homogeneous and heterogeneous spatial domains in our study. The homogeneous domain was set to be a continuous, homogeneous square of size 1 km *×* 1 km. The heterogeneous spatial domain is described subsequently in Section 3.2.1. Analogous to the cervid farms studied in McGregor et al. [2019], Erram et al. [2019], and Erram and Burkett-Cadena [2018], we assume *Culicoides* spp. are able to interact with both domestic and wild hosts, but we only include domestic hosts located on the farm. Hence, midges can fly beyond the limits of the farm domain, however the host population must always remain within the limits. If a midge flies beyond the farm domain, it continues following the same behavior it would as if it were within the domain (Fig. 2).

**Figure 2.**
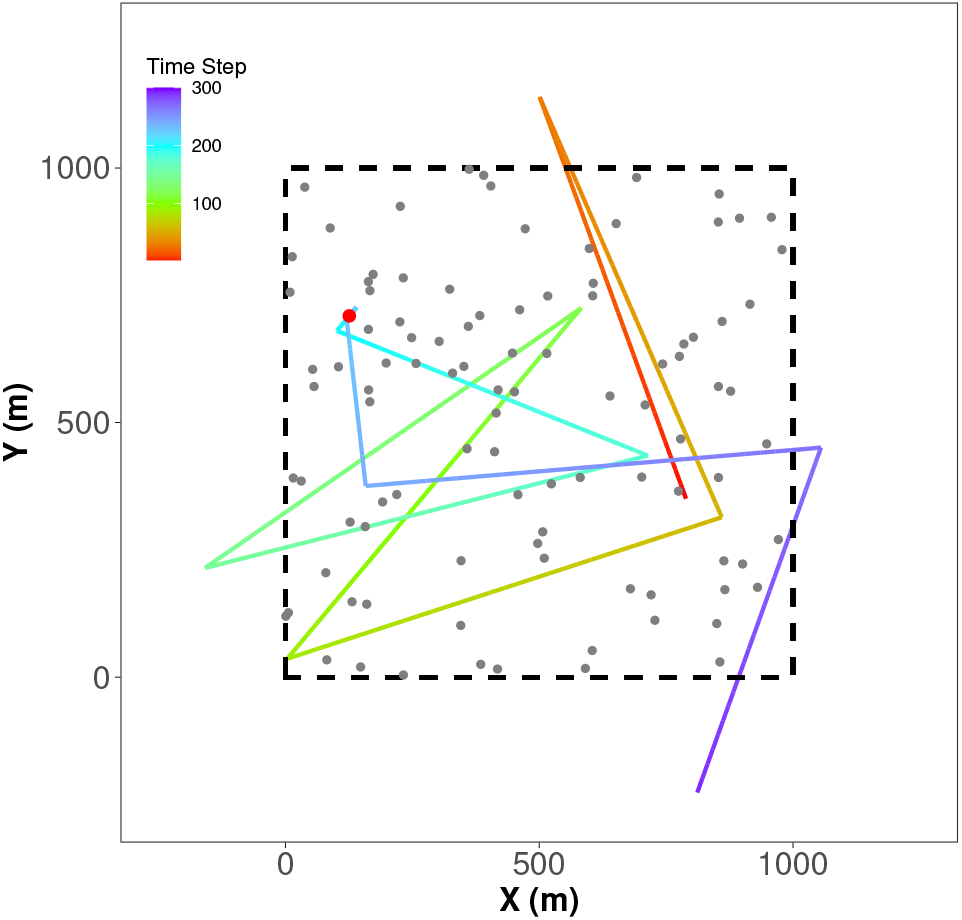
An example path of a midge over one day (300 time steps). The gray dots represent hosts. The red dot indicates the host that the selected midge bit. The dashed boundary indicates the simulated farm domain where the hosts can roam.

At the beginning of each day in the simulation, each ruminant is given a random position within the farm domain. Simulations where ruminants moved in a random walk did not produce notably different results compared to simulations where ruminants remained static throughout each day, and thus, to reduce computational complexity, ruminant locations were updated once daily.

Each day was simulated to be *T*_*d*_ = 5 hours long, which is approximately the amount of time during dawn and dusk where *Culicoides* spp. midges are most active (Fall et al. [2015], González et al. [2017], Blackwell [1997]). At the start of the simulation, an initial number of midges, *I*_0_, are randomly selected to be infected with BTV.

Unless otherwise stated, the simulation period of the model was 60 days. This was chosen as the primary interest for this model was understanding outbreak dynamics, and so longer periods were not necessary.

#### 3.2.1 Heterogeneous Environment

Since midges’ habitat preferences, such as toward bodies of water or swamps, could have an effect on the direction that they would fly (Erram et al. [2019]), we also considered a heterogeneous domain. The domain designed for our study was inspired by the cervid farm located in Gadsden County, FL, USA that was studied in McGregor et al. [2019].

To examine the effects of heterogeneity, we generated a 200 pixel *×* 200 pixel map, where each pixel in the map represents a 5 m × 5 m region in the domain. Each pixel was categorized into one out of five different habitat types, which were given a rank to correspond with the habitat preferences of the midges: 1 is considered the least favorable and 5 is the most favorable.

### 3.3 Midge Movement and Biting Behavior

At each time step, of length Δ*t*, whether a midge has recently fed or not is determined, which then dictates the type of movement that the midge will undergo. We consider a midge to have recently fed if it had a blood meal from a ruminant within the last *T*_full_ days, which we set to be 2 days. The midges that have recently fed (time since last meal *< T*_full_) move in a random walk with roaming flight velocity *v*_*r*_. Letting **x**(*t*) = (*x*(*t*), *y*(*t*)) be the current location of a midge, its location is updated according to

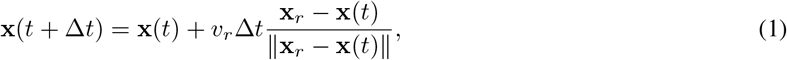

where **x**_*r*_ is a random point uniformly selected within the domain. This ensures that midges remain within the domain and do not travel too far outside the region to detect ruminants when searching for a blood meal.

If a midge has not recently fed (time since last meal ≥*T*_full_), it will check if there is a ruminant within a radius of distance *d*_det_ from itself, which we set to be 300 m. If there are no ruminants within that distance, the midge moves in a random walk with roaming flight velocity *v*_*r*_ as defined above in (1). If ruminants are located within a midge’s detection distance *d*_det_, the midge will fly in the direction of the closest ruminant with active flight velocity *v*_*f*_, such that its location is updated according to

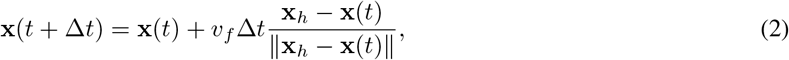

where **x**_*h*_ is the location of the host that is closest to the midge. The midge will move according to (2) until it is within a minimum biting distance *d*_bite_, after which the ruminant will be bitten.

For a heterogeneous domain (Section 3.2.1), we assume that midges undergoing random walk movement would preferentially move toward habitats that are more favorable. Modifications for midge movement are then as follows: at each time step that a midge is not searching for a blood meal, the midge determines the preference ranking of the habitats in its Moore neighborhood (the surrounding eight 5 m *×* 5 m patches as well as the patch that the midge is currently located in). The patches with the highest value according to preference are selected, and then one patch is randomly chosen. The midge flies with roaming flight velocity *v*_*r*_ in the direction of the chosen patch and its location is updated according to

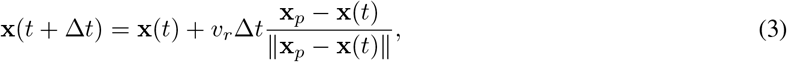

where **x**_*p*_ is the location of the center of the chosen patch (instead of the location being updated according to (1)). The patch selection process is repeated at each time step, so this means that the habitat distribution has an effect on how directed versus how random a midges’ flight movement is: if there is just one patch surrounding the midge with a higher ranking than all of the other patches, the midge will fly directly to that patch. However, if there are multiple patches surrounding the midge with equal preference ranking, then the midge will overall appear to move more in a random walk motion.

### 3.4 Transmission of Bluetongue Virus

After a midge bites a ruminant in the simulation, the subsequent steps are followed: If the host is viremic with BTV, then BTV is transmitted to the midge with probability *P*_HtoV_. If the midge is infectious with BTV, then BTV is transmitted to the host with probability *P*_VtoH_.

After inoculation by a midge, a ruminant that had been infected by BTV would undergo an extrinsic incubation period for *ρ*_H_ days, after which it would become viremic and could potentially infect other midges. Due to a lack of data providing a more precise value, we set *ρ*_H_ to be 2 days. Similarly, a midge that had been infected by BTV would undergo an extrinsic incubation period, *ρ*. The model did not consider the ability of ruminants or midges to recover from BT, and as such they would remain viremic for the duration of the simulation. Furthermore, we do not implement any changes in midge feeding behavior after infection since the biting rate of midges is not well established in the literature and is highly dependent on the data collection and statistical analysis methods employed (Möhlmann et al. [2021]).

Studies have shown that ruminants are often asymptomatic when infected with BT, and can remain viremic for several weeks (Singer et al. [2001]). Therefore due to the short simulation period and the long ruminant lifespan, even with BT, the model did not consider ruminant death.

We assume in each simulation that the midges have a constant daily survival rate, *α*. Laboratory findings have shown that the lifespan of *Culicoides* spp. with respect to a fixed temperature follow either an exponential or Weibull distribution (Lysyk and Danyk 2007]). Assuming an exponential distribution, then the mean expected survival time of the midges is 1*/*(1 − *α*) days. At the end of each day simulated, it is determined whether each midge will survive to the next day with probability *α*. If the midge is determined to have died, it is replaced by a new midge that is not infected with BTV in a random location so that the population size of midges remains constant.

## 4 Results

In this section, we analyze the sensitivity of the model parameters to the model output (Section 4.1), determine the dependence of the number of infected ruminants on *Culicoides* spp. survival rate, extrinsic incubation period, and temperature using midge flight activity as a proxy (Section 4.2), evaluate the dependence of outbreak probability on the initial number of infected midges, their survival rate, and probability of transmission from vector to host (Section 4.3), and examine the effects of a heterogeneous environment on midge location preferences (Section 4.4).

### 4.1 Sensitivity Analysis

We determine how sensitive the model is to changes in the *Culicoides* spp. daily survival rate, *α*, extrinsic incubation period, *ρ*, probability of BTV transmission from vector to host *P*_VtoH_, probability of BTV transmission from host to vector *P*_HtoV_, and initial number of infected midges, *I*_0_, using Sobol’ sensitivity analysis (Sobol’ [2001]) via Saltelli’s extension of the Sobol’ sequence (Saltelli [2002], Saltelli et al. [2010]) as implemented in the SALib Sensitivity Analysis Library (Herman and Usher [2017]). This is a variance-based method which allows for calculation of model output changes with respect to variation with a single parameter (first-order), and combinations of all parameters (total-order). By comparing the first-order and total-order indices, the presence of higher-order interactions can be inferred.

The range of values for the parameters of interest were set to be *α* ∈ [0, 1], *ρ* ∈ [0, 20], *P*_VtoH_ ∈ [0, 1], and *P*_HtoV_ ∈ [0, 1], which contain the typical values observed for these parameters (Table 1). The total number of infected ruminants after 60 days was used as the model output, and the resulting first- and total-order indices were calculated. Due to computational constraints and the stochastic nature of the model, a total of 96 simulations, with parameters given by the Saltelli distribution, were run for each trial.

To clarify terminology used in this section, we define a simulation to be a single execution of the model under a single set of parameters. A trial is a group of simulations. We averaged the results from many trials to find the expected values for the analysis in this section.

Both the first-order indices and total-order indices, shown in Fig. 3, indicated that the model was much more sensitive to perturbations in the *Culicoides* spp. survival rate *α* than it was to perturbations in the extrinsic incubation period *ρ* or the transmission probabilities *P*_VtoH_ and *P*_HtoV_. Thus, this analysis implies that focusing on the survival of midges is more important than the incubation period and the probability of BTV transmission from host to vector or vice versa in reducing the number of infected ruminants.

**Figure 3.**
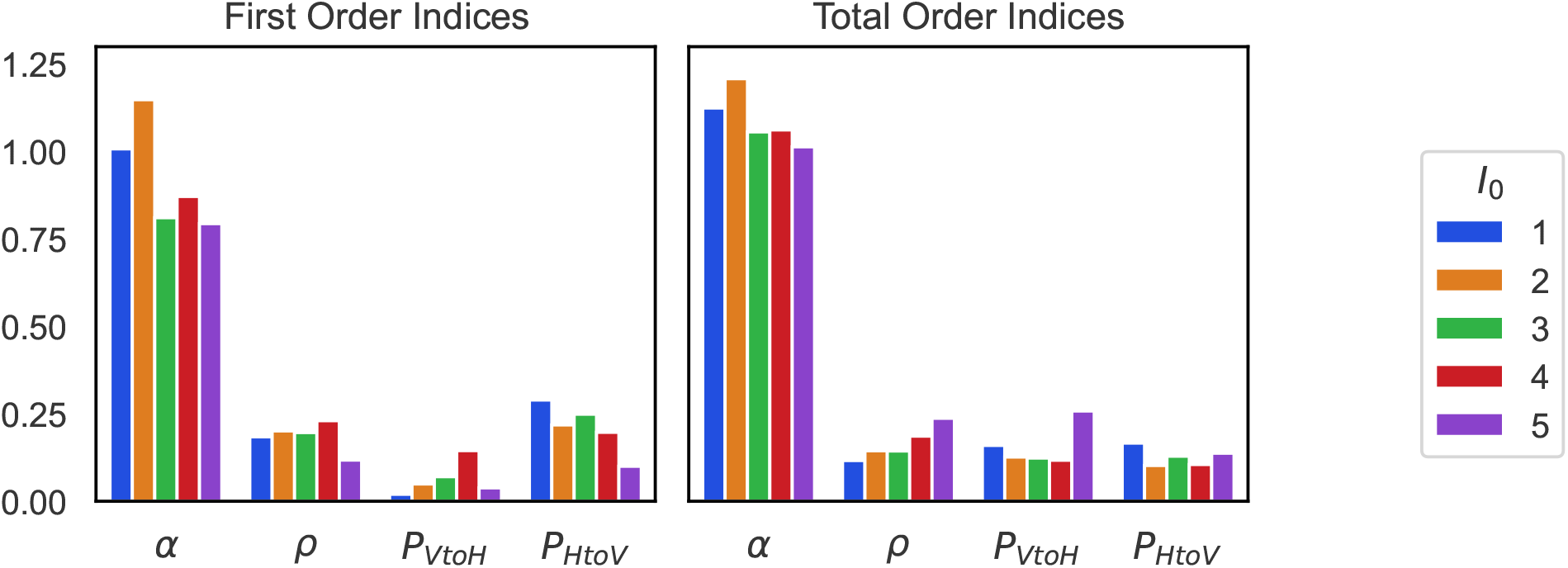
First-order and total-order indices for Sobol’ sensitivity analysis of the *Culicoides* spp. daily survival rate, *α*, the extrinsic incubation period, *ρ*, the probability of BTV transmission from vector to host *P*_VtoH_, and the probability of transmission from host to vector *P*_HtoV_. *I*_0_ indicates the initial number of BTV infected midges. First-order indices are calculated with respect to a single parameter, so interactions with other parameters are not taken into account, while total-order indices account for all higher-order interactions between parameters.

Additionally, it was found that the model sensitivity to *α* decreased as *I*_0_ increased, which was to be expected as more BTV-infected *Culicoides* spp. midges would reduce the need for high survival rates. Initial analysis was conducted using 10 trials for each *I*_0_, and then was increased to 15 trials. Little change was noticed, and so the analysis shown was conducted using 15 trials.

### 4.2 Percentage of Infected Ruminants

A heat map, shown in Fig. 4, was generated to visualize the total percentage of infected ruminants over a 60 day period given changes in the *Culicoides* spp. survival rate *α* and extrinsic incubation rate *ρ*. We also varied the total midge population, as given by the number of midges per ruminant, *γ*. Since midges are more active in warmer temperatures (Tugwell et al. [2021], increasing *γ* implies that there is more midge flight activity, which is correlated with warmer temperatures. Hence, we can implicitly examine the effects of temperature by varying *γ* as a proxy. Parameter values were chosen to be *α* ∈ [0.6, 0.9] in increments of 0.015, *ρ* ∈ [10, 20] in increments of 0.5, and *γ* ∈ {5, 10, 50, 100}. For each combination of parameters *α, ρ*, and *γ*, 10 simulations of each case were run due to the stochastic nature of the model and the tendency for low values of *α* or high values of *ρ* to result in no outbreak at all. The average number of infected ruminants was averaged over all the simulations for each (*α, ρ, γ*)-set.

**Figure 4.**
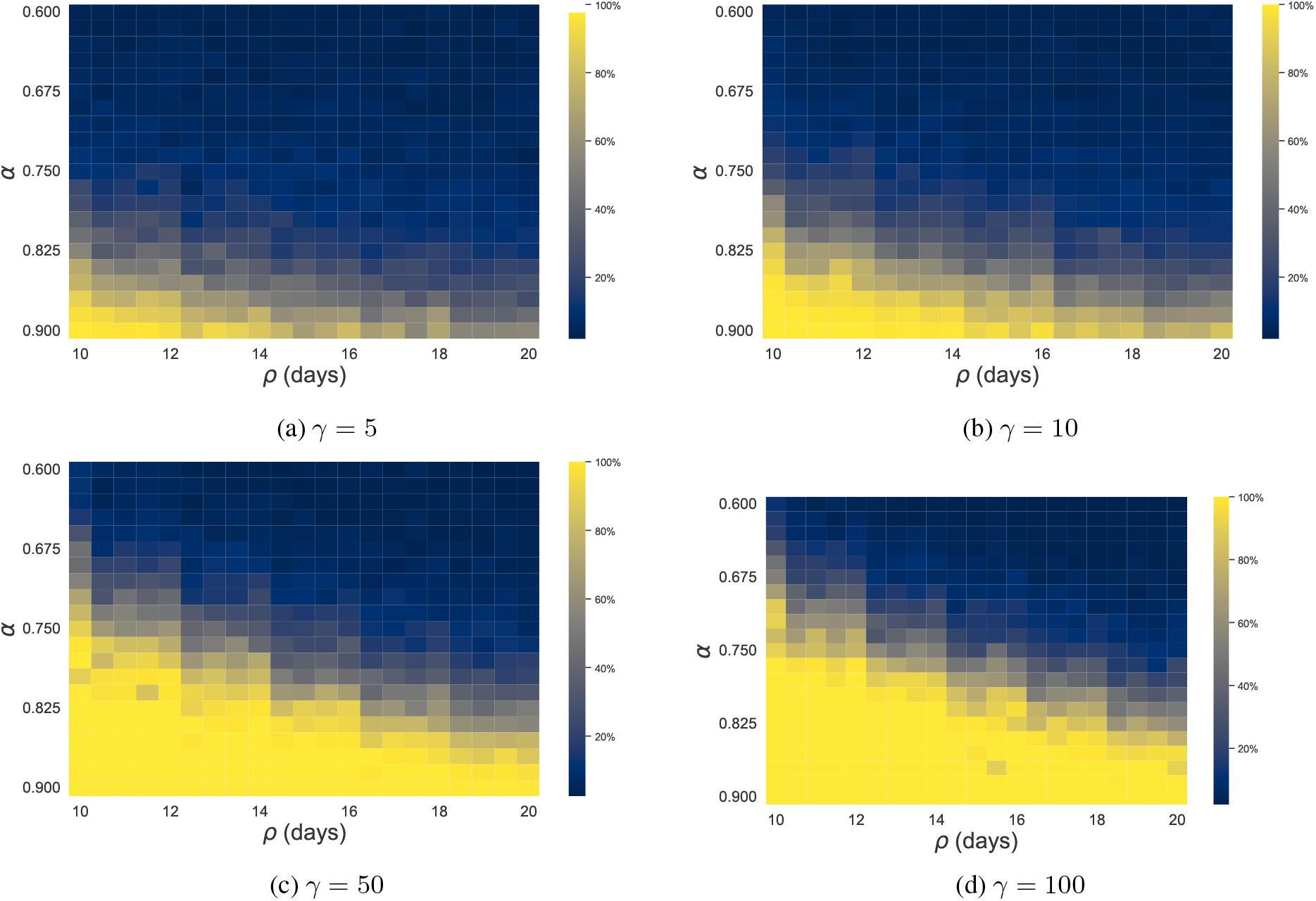
Heat maps showing the percentage of ruminants infected given different *Culicoides* spp. daily survival rate, *α*, extrinsic incubation rate, *ρ*, and number of midges per ruminant, *γ*, over a 60 day period.

Fig. 4 illustrates that the number of infected ruminants is non-decreasing as *α* and *ρ* increase for each fixed value of *γ*, and it was very common for simulations with large enough values of *α* and *ρ* to end with all of the ruminants within the domain being infected with BTV. There appears to be a linear bifurcation line with respect to *α* and *ρ* that determines the likelihood of an outbreak. Low values of *α* combined with high values of *ρ* would result in few to no outbreaks, while high *α* and low *ρ* result in large-scale outbreaks. This line shifts as *γ* increases, such that large midge populations correspond to larger outbreaks for similar combinations of *α* and *ρ*. This follows intuition, as a high survival rate and low extrinsic incubation period would allow many more *Culicoides* spp. midges to become infected with BTV and subsequently pass the virus on to another ruminant. Additionally, in warmer temperatures where midge flight activity is greater, there is an increased chance of an outbreak.

### 4.3 Outbreak Probability

For the purposes of this study, an outbreak was defined to be the occurrence of a single BTV-infected ruminant. We fixed the initial number of infected midges, *I*_0_, and daily survival rate, *α*, for each simulation and allowed the simulation to run until either all BTV-infected midges died without a single ruminant becoming infected or a single ruminant had become infected with BTV. Parameter values were chosen to be *α* ∈ [0, 1] in increments of 0.02, *P*_VtoH_ ∈ {0.25, 0.5, 0.75, 1}, and *I*_0_ ∈ {1, 2, 3, 4, 5, 15, 50, 100}. 500 simulations were run for each (*α, P*_VtoH_, *I*_0_)-set. From this, the estimated outbreak probability was calculated by averaging over all the simulations.

Fig. 5 illustrates that for low values of *I*_0_, the relationship between *α* and the probability of outbreak is concave up for small *P*_VtoH_ and then is approximately linear for large *P*_VtoH_. As *I*_0_ increases, especially past *I*_0_ = 15 and for any value of *P*_VtoH_, the outbreak curves approach a step function. As *P*_VtoH_ increases but *I*_0_ is kept fixed, the concavity of the outbreak curves decreases, i.e., a concave up curve becomes more linear and a concave down curve approaches a step function. This indicates that an increase in *α, I*_0_, or *P*_VtoH_ corresponds with higher outbreak probabilities.

**Figure 5.**
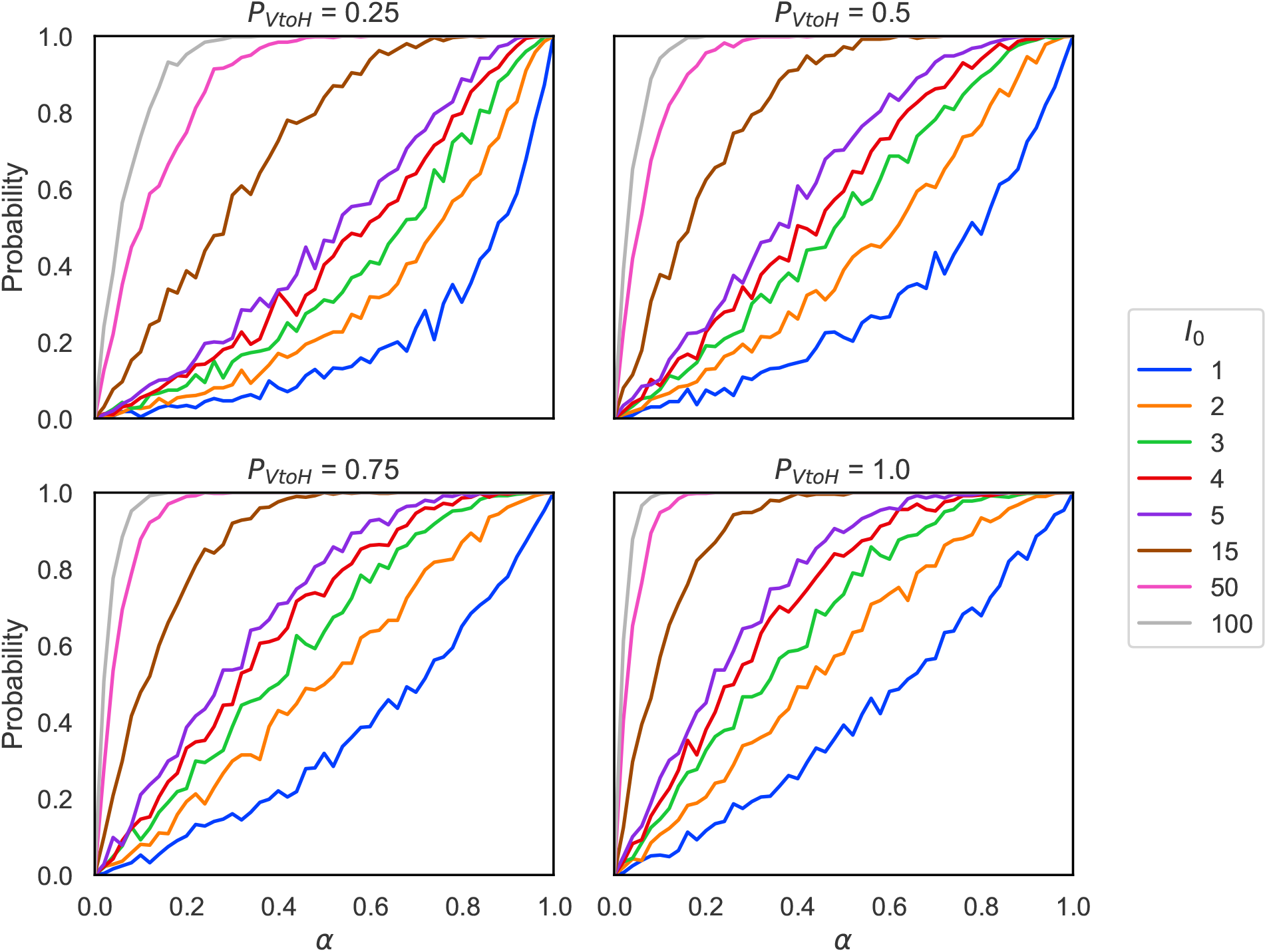
The probability of outbreak given different initial numbers of infected midges, *I*_0_, depending on the daily survival rate, *α*, and the probability of transmission from vector to host *P*_VtoH_. An outbreak was defined to be the infection of a single ruminant. The probability of outbreak increases dramatically as *α* increases. The shape of the curve is concave up or linear for low *I*_0_ and for low *P*_VtoH_ or high *P*_VtoH_, respectively, and approaches a step function for high *I*_0_, regardless of the value of *P*_VtoH_.

### 4.4 Heterogeneous Environment Simulation

The effects of a heterogeneous environment are studied using the map illustrated in Fig. 6a, which was inspired by the cervid farm studied in McGregor et al. [2019]. Following the additional movement rules as described in Section 3.3, midges preferentially travel to habitats that are more favorable (3) instead of random walk motion (1). Due to computational constraints, a single simulation of 30 days was run, and midge locations were saved at the final time step. The spatial density of the midges is shown in Fig. 6b.

**Figure 6.**
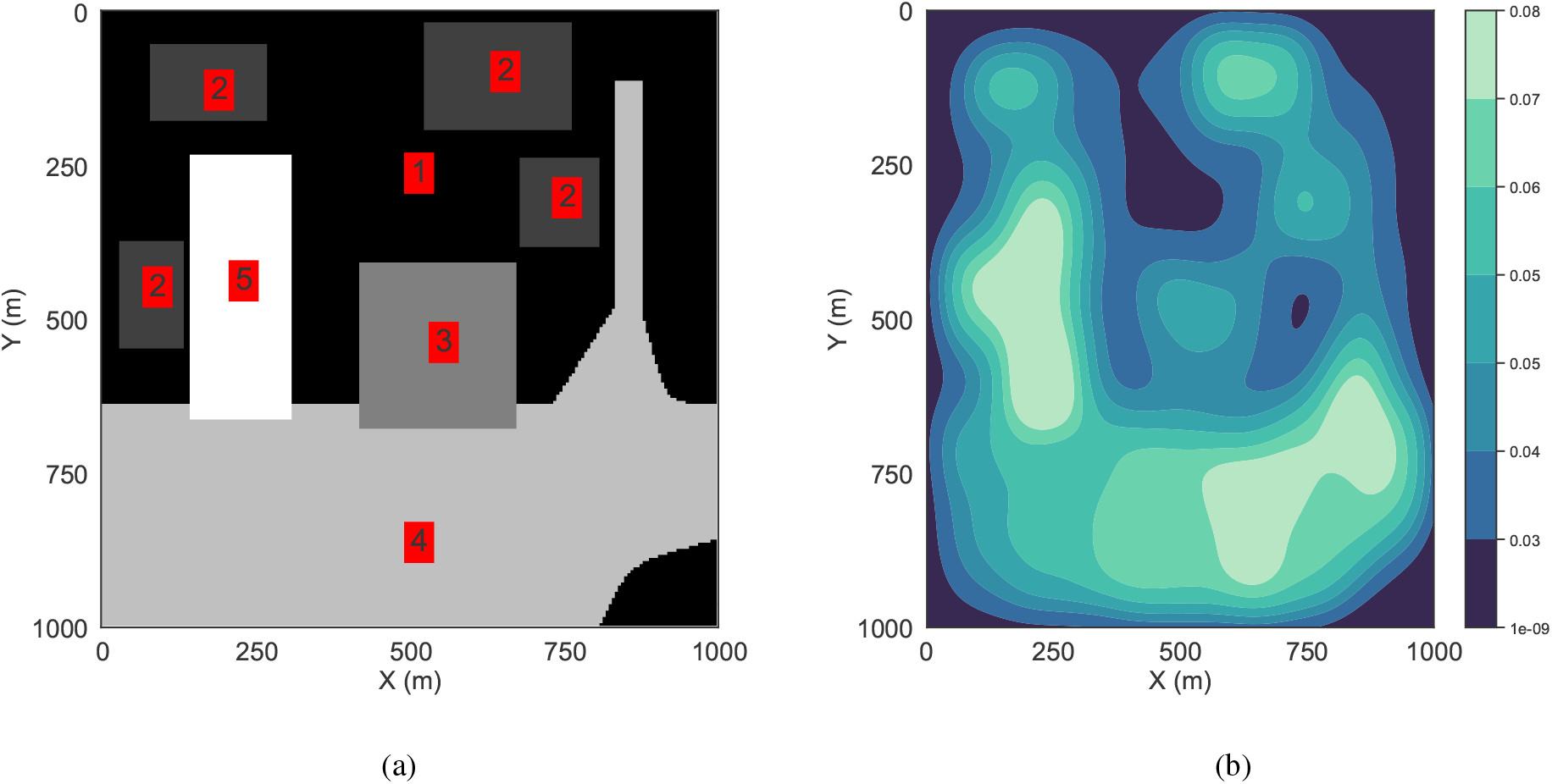
Heterogeneous environment simulation. (a) Map of a heterogeneous domain, where lighter colors and higher numbers indicate habitats that are more favorable to midges. (b) A density plot of simulated midges at 30 days in the heterogeneous environment, with *α* = 0.75, *P*_*d*_ = 100, and *γ* = 500. Units: midges per m^2^.

Fig. 6 indicates that there are strong structural similarities between the density distribution of the simulated midges (Fig. 6b) and the preferred regions in the map (Fig. 6a). Specifically, there is a long vertical cluster of midges centered near (200, 500), which corresponds to the most preferred region in the map (level 5). An additional highly-dense cluster is centered near (700, 800), which corresponds to the next most preferred region of level 4, observing that this cluster is closer to regions of level 3 and 1, rather than the level 5 region. Finally, there is a noticeable lack of midges near (500, 250), which is linked to the large and least preferred region of level 1.

## 5 Discussion

The goal of this paper was to develop a model that represents the spread of BTV in a controlled environment. This model was investigated through methods that included sensitivity analysis and an examination of the probability of outbreak. Results indicated that the model was more sensitive to variations in *Culicoides* spp. survival rate than to the extrinsic incubation period of BTV or transmission probability (Fig. 3), which was reinforced by analysis of the probability of outbreak (Fig. 5). Additional visual analysis of the heat map in Fig. 4 suggests that there exists a bifurcation when considering the relationship between survival rate and extrinsic incubation period, which specifies parameter regimes in which outbreaks are expected. As such, we conclude from this model that there exists a strong relationship between survival rate and the spread of BTV.

It is first worth noting the limitations of the model. As it is extremely difficult to accurately count the population of biting midges in a given system, the population of midges was set to be 100 times the number of ruminants. This ratio was chosen due to computational constraints. It is very likely that this is an underestimation of the total number of midges, and so the true spread of BTV would occur among a much larger vector population. Additionally, the model does not consider ruminants that ultimately recover or die from BTV infection, thereby resulting in a non-decreasing number of viremic hosts that will indefinitely contribute to the spread of BTV. Therefore it is possible that the model is overestimating the number of infected ruminants over long simulation periods (greater than one or two months).

Simulations of the model suggest that early in the outbreak, BTV could be in the system while not being detected. This would occur during a short window where there are no viremic ruminants or midges capable of spreading BTV, and instead, the virus is within a host animal or midge after inoculation has occurred but the virus has not yet completed its replication cycle. This suggests that it is much more difficult for ecologists and epidemiologists to detect and isolate early outbreaks of BTV in a region. We suggest that it would be possible to counter this issue by repeated testing of susceptible hosts over multiple days in at-risk areas.

A key focus of this model was analyzing possible outbreak scenarios. As such, the analysis has shown that survival rate of the midges plays the most important factor in determining the probability of an outbreak. Other studies have also shown that vector mortality rate and density significantly affect disease spread within host populations for mosquitoes transmitting malaria (Mandal et al. [2011]) and tsetse flies transmitting human African trypanosomiasis (Gervas et al. [2018]). Therefore it would be prudent that, in regions at risk of an outbreak, the main focus should be to reduce the survival rate of vector species through means such as pest control. Other options include temporary removal or isolation of BTV host species, which most commonly are livestock used in agriculture.

One result of the model is that the outbreak probability increases with *I*_0_ (Fig. 5). While the trend appears linear for low values of *I*_0_, it was found that this trend becomes increasingly concave. It appears that in the limit as *I*_0_ increases, the probability of outbreak would approach a step function. While this follows intuition, and helps to further validate the model, the results further suggest that high numbers of infected *Culicoides* spp. midges virtually guarantee the possibility of an outbreak. This implies that population control measures such as pesticides may not be enough to prevent epizootics, as long as there are enough vectors already infected with BTV to continue the cycle. This presents a predicament, as another strategy to prevent outbreaks is mass vaccination of livestock in the surrounding region. However, the development of new vaccines is challenging due to the increasing appearance of new BTV serotypes which are not immunologically simple (Maclachlan et al. [2015]), increasing the economic impact through very expensive vaccine development and administration (Gerry et al. [2001]). Ideally, outbreaks would be controlled by a combination of population-control methods and vaccination programs.

Multiple studies have shown a strong seasonality with the growth and decay of BTV (Gerry et al. [2001], Carpenter et al. [2011]). It is known that BTV can lie dormant for multiple months during colder, more unfavorable conditions, then return during spring and summer where its prevalence reaches its peak. This model did not attempt to simulate outbreaks on large time scales, and so seasonality was not considered in the design. Instead, the model assumes constant parameters and consistent environmental variables. Additionally, one of the key goals of this model was to understand BTV spread in the southeast United States, particularly North Florida, where BTV remains endemic and does not spread through outbreaks like in Europe. In the southeast and North Florida in particular, the subtropical environment offers less seasonality than in more northern regions. As such, the lack of seasonality in this model is not as critical of a limitation as it would be in a model for more temperate regions.

In simulations with a heterogeneous environment where midges follow preferential movement toward more favorable habitats, the midges showed a strong ability to arrange themselves in habitats that are more preferable, such as those commonly being bodies of water or swamps (Erram et al. [2019]). Furthermore, as illustrated in Fig. 6b, midges will not only concentrate in the highest favored habitats, as expected, but there can be less favorable habitats that midges will also cluster in, such as the level 4 region centered around (700, 800) which is closer to level 1 regions than expected. Thus, regions containing high midge concentrations using MidgePy simulations can be used to suggest optimal farm layouts that involve organizing pastures and fences to avoid these areas. As Fig. 4 indicates, smaller midge populations correspond to fewer BTV infected ruminants for similar *ρ* and *α*. Therefore habitats which are suitable for ruminants and not preferable for midges should be prioritized to reduce BT outbreak occurrence and severity.

Future work will focus on the development of a continuous model based on differential equations to identify a functional relationship between the *Culoicoides* spp. daily survival rate *α* and extrinsic incubation period *ρ*. While the sensitivity analysis indicates *α* to be more influential than *ρ* in determining the size of an outbreak, *ρ* still plays a significant role. In fact, there appears to be a linear relationship between *α* and *ρ* that shows how high survival rates may not be enough to guarantee an outbreak when combined with long incubation periods.

This model was designed with simplicity in mind. For that reason, it would be straightforward to modify the model in order to investigate the spread of other vector-borne diseases. Other future avenues of research using this model as a base include modeling malaria, which is spread through humans and other animal hosts. This would add a new level of complexity as humans move significant distances more frequently than livestock. Diseases such as malaria could be implemented into the model with relative ease, and sensitivity analysis could be performed to understand better methods for controlling spread. Additionally, there are other model parameters which are of potential future interest, including the incubation period of BT within ruminants, *ρ*_H_, and midge biting rate, *T*_full_. These parameters warrant further study, as well as extending the model to include vaccination and ruminant death to account for the high mortality rate found in some ruminants, particularly sheep, due to BTV (Conraths et al. [2009]). Such analysis could supplement biological research on BTV-host interactions.

## Supplementary information

The source code for MidgePy used to generate the results for this article is available through GitHub at https://github.com/stepien-lab/MidgePy [v1.0.0]. The code is platform independent and written in Python. The data used for this article is available through OSF at https://www.osf.io/fhven.

## Acknowledgments

The authors acknowledge University of Florida Research Computing for providing computational resources and support that have contributed to the research results reported in this publication.

T.L.S. acknowledges support from a Simons Collaboration Grant for Mathematicians (#710482) and NSF grant DMS-2151566.

## Statements and Declarations

### Competing interests

The authors declare no competing interests.

## Appendices

## Notes

### Competing Interest Statement

The authors have declared no competing interest.

### Summary of Updates

Updated results to include expanded parameter analysis and examine the effects of a heterogeneous environment.

